# A single-cell atlas of the human healthy airways

**DOI:** 10.1101/2019.12.21.884759

**Authors:** Marie Deprez, Laure-Emmanuelle Zaragosi, Marin Truchi, Sandra Ruiz Garcia, Marie-Jeanne Arguel, Kevin Lebrigand, Agnès Paquet, Dana Pee’r, Charles-Hugo Marquette, Sylvie Leroy, Pascal Barbry

## Abstract

**Rationale:** The respiratory tract constitutes an elaborated line of defense based on a unique cellular ecosystem. Single-cell profiling methods enable the investigation of cell population distributions and transcriptional changes along the airways.

**Methods:** We have explored cellular heterogeneity of the human airway epithelium in 10 healthy living volunteers by single-cell RNA profiling. 77,969 cells were collected by bronchoscopy at 35 distinct locations, from the nose to the 12^th^ division of the airway tree.

**Results:** The resulting atlas is composed of a high percentage of epithelial cells (89.1%), but also immune (6.2%) and stromal (4.7%) cells with peculiar cellular proportions in different sites of the airways. It reveals differential gene expression between identical cell types (suprabasal, secretory, and multiciliated cells) from the nose (*MUC4*, *PI3*, *SIX3*) and tracheobronchial (*SCGB1A1*, *TFF3*) airways. By contrast, cell-type specific gene expression was stable across all tracheobronchial samples. Our atlas improves the description of ionocytes, pulmonary neuro-endocrine (PNEC) and brush cells, which are likely derived from a common population of precursor cells. We also report a population of *KRT13* positive cells with a high percentage of dividing cells which are reminiscent of “hillock” cells previously described in mouse.

**Conclusions:** Robust characterization of this unprecedented large single-cell cohort establishes an important resource for future investigations. The precise description of the continuum existing from nasal epithelium to successive divisions of lung airways and the stable gene expression profile of these regions better defines conditions under which relevant tracheobronchial proxies of human respiratory diseases can be developed.

## Introduction

The prevalence of chronic respiratory diseases is thought to increase in the future by raising exposure to diverse atmospheric contaminants (pollution, allergens, smoking). The respiratory tract constitutes an elaborated line of defense based on a unique cellular ecosystem. Thus, secretory and multiciliated cells form a self-clearing mechanism that efficiently removes inhaled particles from upper airways, impeding their transfer to deeper lung zones. Several mechanical filters (nose, pharynx, ramified structure of the lung airways) further limit the flux of pathogens and inhaled particles downwards the bronchial tree. While nose and bronchus are sharing many cellular properties, which has led to the definition of a pathophysiological continuum in allergic respiratory diseases (1, 2), they differ by features such as host defense against viruses, oxidative stress (3), or anti-bacterial mechanisms (4). Bulk RNA sequencing has indeed identified differences in gene expression between nose and bronchi (5). The recent advent of single-cell RNA sequencing offers an excellent opportunity to carefully analyze and compare cellular composition and gene expression from nasal to the successive generations of the lung airways. In the framework of the Human Cell Atlas (HCA) consortium, we have now established a precise airway epithelium cell atlas in a population of 10 healthy living volunteers. Minimally invasive methods were set up to collect biopsies and brushings using bronchoscopy. A high-quality dataset of 77,969 single cells comprising a large panel of epithelial cell subtypes was generated from 35 distinct samples taken at precise positions in the nose, trachea and bronchi. Data integration and analysis provide a unique view of the cell type proportions and gene signatures from the first to approximately the 12^th^ division of the airways. The resulting picture defines a relatively stable cellular composition and gene expression across the first 12 successive generations of the tracheobronchial tree. The largest differences were found between nasal and bronchial samples. Our work better defines the conditions under which the nose can be considered as a relevant bronchial surrogate for studying human lung pathologies, and underlines the relative stability of gene expression in the proximal tracheobronchial compartment of the airways.

## Methods

The atlas of the airway epithelium (nose to approximately 12^th^ division of bronchi) was obtained from biopsies and brushings from 10 healthy non-smoking volunteers. Each donor was sampled 4-5 times in different regions of upper (nose) and lower airways (tracheal, intermediate, distal bronchi), located in different lobes (Figure E1, Tables E1-2). Single-cell capture was carried out using the 10X Genomics Chromium device (3’ V2). Large integrative analysis of the 35 samples composing the atlas was done using the fastMNN R package (6) and the following analysis was done using the Scanpy framework (Python) (7) to provide robust cell type annotation. Additional differential gene expression analysis was done using the edgeR R package (8) to investigate both cell distributions and gene expression heterogeneity along the airways. GSEA, trajectory inference (PAGA) (7) and gene network inference (GRNBoost2) (7) were also performed to characterize further the identified cell populations. Results were validated using RNAscope and immunostainings. Additional details on the methods are provided in an online data supplement.

## Results

### Building a molecular cell atlas of the airways in healthy volunteers

#### Data collection

Cells were analyzed by droplet-based single-cell RNA sequencing (scRNA-seq), after isolation from 4 distinct locations using 2 sampling methods: (i) nasal biopsies (3 samples) and (ii) nasal brushings (4 samples), (iii) tracheal biopsies (carina, 1^st^ division, 9 samples), (iv) intermediate bronchial biopsies (5-6^th^ divisions, 10 samples), (v) distal brushings (9-12^th^ divisions, 9 samples) in 10 healthy volunteers (Figure 1A, 1B, Figure E1A, Table E1). Optimized handling and dissociation protocols allowed the profiling of 77,969 single cells which were collected at 35 distinct positions of the airways, resulting in the detection of an average of 1,892 expressed genes per cell with 7,070 UMI per cell (Figure E2A).

**Figure 1.**
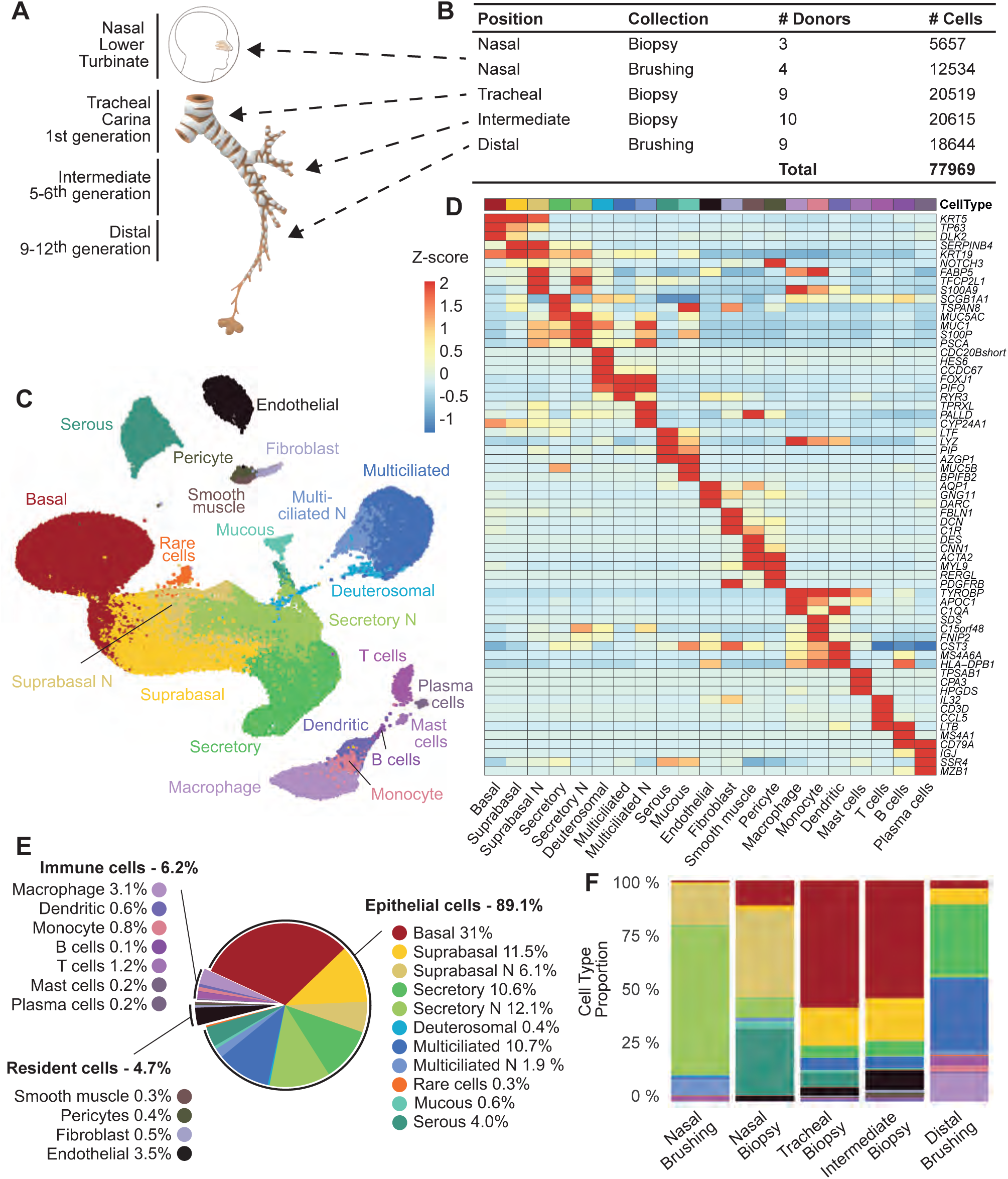
A molecular cell atlas of the healthy human airways. **(A)** Schematic representation of the sampled anatomical regions. **(B)** Experimental design of the study, detailing the anatomical regions, sampling methods, number of donors, biopsies and cells after data curation. **(C)** UMAP visualization of the whole human healthy airway dataset. Each distinct cell type is defined by a specific color. **(D)** Heatmap of expression for top marker genes of each cell type. **(E)** Pie chart of the total proportion of each cell type identified in human airways. **(F)** Barplot of the relative abundance of each cell type collected by two distinct modes of biopsies at four macro-anatomical locations.

Following batch correction and graph-based clustering, cell types were assigned to each cluster using well-established sets of marker genes (Figure 1C, Figure E3). We identified 14 epithelial cell types, including 12 for the surface epithelium and 2 for submucosal glands, which collectively represented 89.1% of total cells (Figure 1C-1E, Table E2; See also our interactive web tool https://www.genomique.eu/cellbrowser/HCA/?ds=HCA_airway_epithelium). Stromal and immune cells represented respectively 4.7% and 6.2% of all cells (Figure 1E).

#### Annotation of epithelial cells

Basal cells (*KRT5*, *TP63* and *DLK2*-high) accounted for one-third of all cells (Figure 1D and 1E). We also identified suprabasal cells, characterized by low *TP63* expression, decreasing gradients of *KRT5* expression and increasing gradients of *KRT19* and *NOTCH3* expression (9–12) (Figure 1D). We grouped club and goblet cells as “secretory cells” since these two populations could not be clustered separately and essentially differed by the level of expression of *MUC5AC* and *MUC5B* (Figure E4) (12). We detected clusters of multiciliated cells (expressing high levels of *FOXJ1*, *TPPP3*, and *SNTN*) and deuterosomal cells, which correspond to precursors of multiciliated cells and express several specific markers: *CCDC67*/*DEUP1*, *FOXN4* and *CDC20B* (Figure 1C and 1D) (12, 13). The suprabasal, secretory and multiciliated clusters each comprised a sub-cluster of cells that were only detected in nasal samples. These clusters were labelled “Suprabasal N”, “Secretory N” and “Multiciliated N” and will be described later in the manuscript. Two cell types were associated with submucosal glands: serous cells (expressing high levels of *LTF*, *LYZ* and *PIP*) and mucous cells (expressing high levels of *MUC5B* but no *MUC5AC*) (Figure 1C and 1D). Finally, we identified 222 cells belonging to clusters of rare epithelial cells (0.3% of the cells) (Figure 1C and 1D). Incidentally, we detected the presence of some alveolar cells: 10 type I (AT1) and 11 type II (AT2) pneumocytes, which were all derived from a unique distal brushing. AT1 expressed *HOPX*, *AGER*, *SPOCK2*; AT2 expressed *SFTPA*, *SFTPB*, *SFTPC* and *SFTPD* (Figure 3B, Table E2, Figure E5).

**Figure 2.**
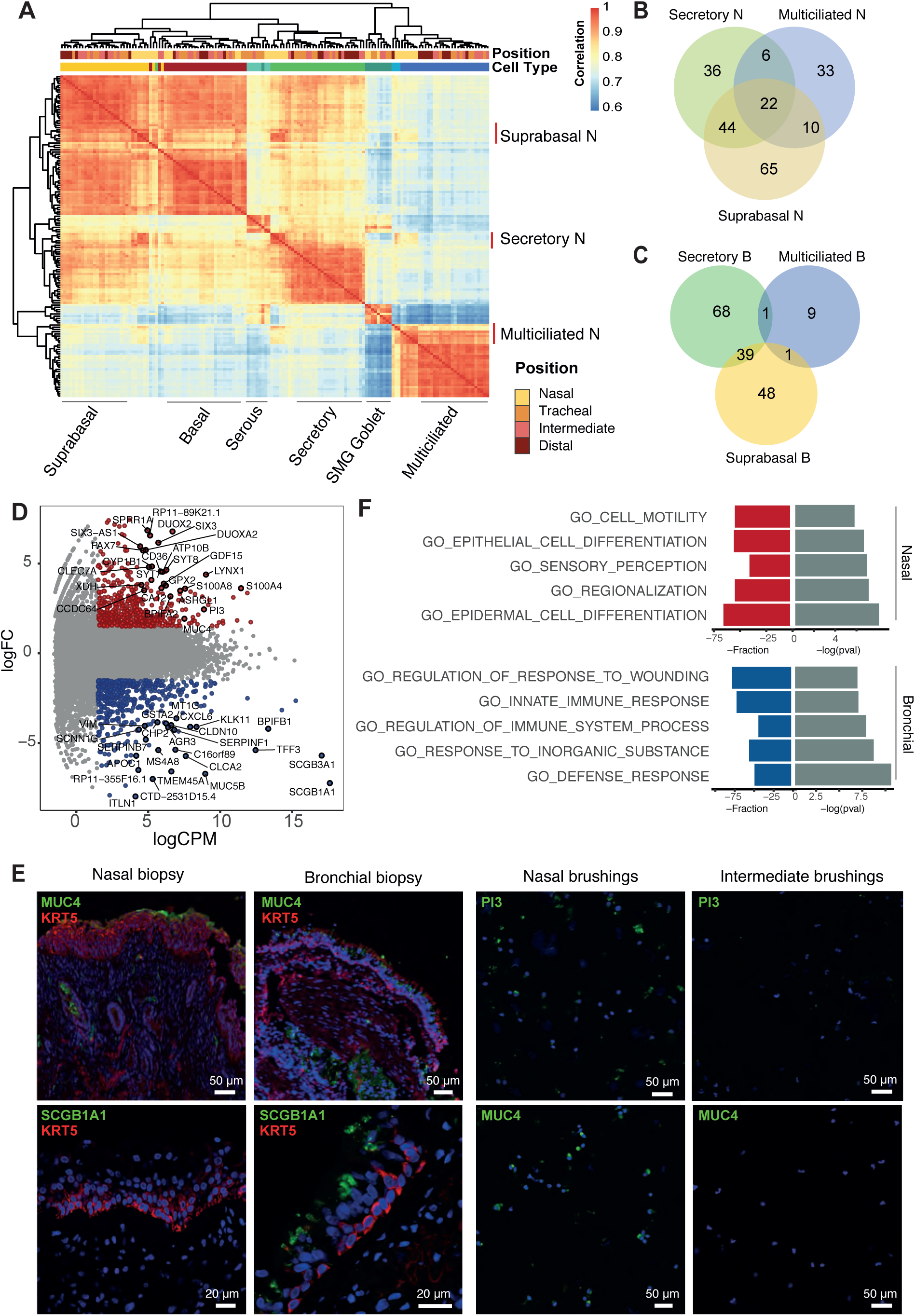
Distinct gene expression signatures are detected between nasal and tracheobronchial airways. **(A)** Unsupervised hierarchical clustering of gene expression correlation between sample-specific cell types. **(B-C)** Venn Diagrams indicating the number of specific transcripts of each cell type (secretory, suprabasal and multiciliated cells), in the nose **(B)** and tracheobronchial airways **(C)**. **(D)** MA-plot of differential expression between secretory cells from the nose versus tracheobronchial airways. Red and blue dots indicate nasal and tracheobronchial airways over-expressed genes, respectively. **(E)** Detection by immunofluorescence of proteins that are more specifically associated with a nose or a tracheobronchial expression. **(F)** Enriched gene sets associated with nasal and tracheobronchial secretory cell markers.

**Figure 3.**
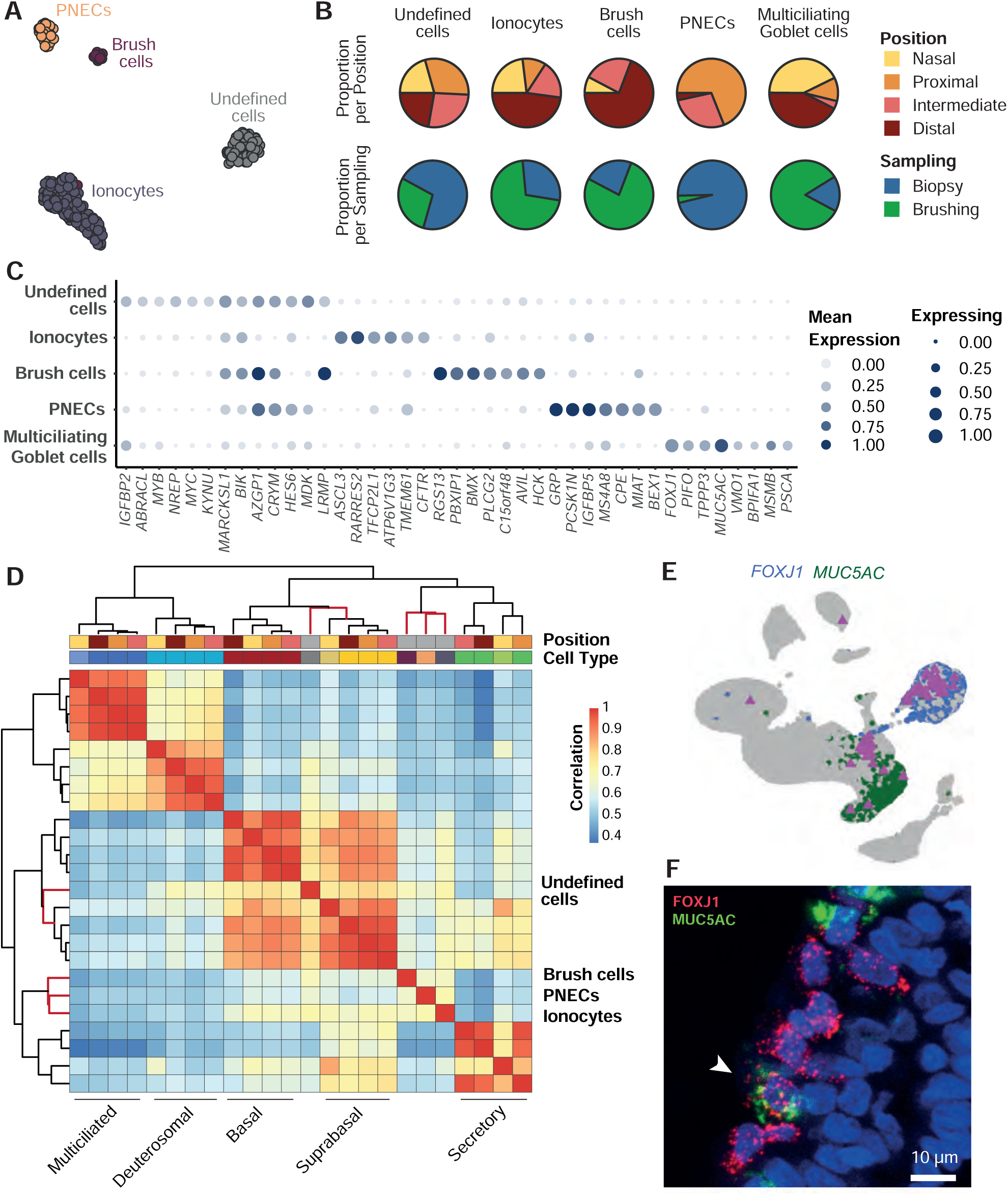
Detection of rare epithelial cells across human airways. **(A)** Focused UMAP visualization on the group of ionocytes, neuroendocrine, brush cells and undefined cells. **(B)** Pie charts of the anatomical distribution of each cell type according to location (top line) or mode of sampling (bottom line). **(C)** Dot plot of the top gene markers identified per cell type of interest. **(D)** Unsupervised hierarchical clustering of gene expression correlation between position-specific epithelial cell types. **(E)** UMAP visualization of double positive *FOXJ1+-MUC5AC+* cells (purple), relative to *FOXJ1+* cells (blue) and *MUC5AC+* cells (green). **(F)** RNAscope detection of a mucous-multiciliated cell in nasal tissue. Red: *FOXJ1+* RNA; green: *MUC5AC+* RNA and white: *SCGB1A1+* RNA.

#### Immune cells: annotation and distribution along the respiratory tree

We clustered the 4891 immune cells in 7 distinct cell types (Figure 1C-1E, Figure E6A). Four clusters of myeloid cells were found: (i) macrophages and (ii) monocytes, mostly detected in distal brushings; (iii) mast cells, mostly detected in distal brushing as well and at a lesser extent in tracheal and intermediate bronchial biopsies; (iv) dendritic cells, found everywhere. We also identified 3 clusters of lymphoid cells: T cells were found in all samples; plasma cells were exclusively found in biopsies, in line with an interstitial localization and B cells were mostly detected in distal airway brushings (Figure 1F, Figure E1B-D). The gene regulatory network was further characterized with GRNboost2, a program that infers regulatory unit activity (14) (Figure E6B). In the lymphoid lineage, we were able to discriminate B cells (expressing high levels of *MS4A1* and *LTB*, and high *PAX5* inferred activity) from plasma cells (expressing high levels of *IGJ* and *MZBI*, and high *IRF4* inferred activity) (Figure 1D, Figure E6A, B). T cells and related subtypes, that our analysis did not separate well, were characterized by a high and specific transcriptional activity of the *XCL1* and *CD3D* regulatory units (Figure E6B, E6C and E6E).

#### Stromal cells: annotation and distribution along the respiratory tree

Stromal cells were found only in biopsies, especially in intermediate samples (Figure 1F, Figures E1B and E7B). We annotated 4 stromal cell types (Figure 1C-1E) including endothelial cells, expressing high levels of *DARC*, fibroblasts, expressing high levels of *FBLN1*, as well as smooth muscle cells, characterized by high levels of desmin (*DES*) and high activity of the *HOXA4* regulatory unit (Figure 1D, Figure E7C). Based on a specific expression of markers such as *RERGL*, *MCAM* and *PDGFRB*, we also identified pericytes, a population of peri-endothelial mesenchymal cells with contractile properties that are located on the vascular basement membrane of capillaries (15, 16). Pericytes also share with smooth muscle cells markers such as *ACTA2* and *MYL9* (Figure 1D, Figure E7D).

### Large variations in the composition of epithelial cells distinguish nasal and tracheobronchial airways

We then compared the epithelial composition in each of the 5 types of samples. We noticed a large effect produced by the sampling mode on the distribution of cells: brushings collected more luminal cell types, such as multiciliated or secretory cells, while forceps biopsies collected cells located deeper in the tissue such as basal, stromal cells, and submucosal gland cells (Figure 1F, Figure E1B, Table E2). All following comparisons were then performed on samples obtained with similar sampling methods.

Tracheal and intermediate bronchial biopsies shared very similar cell type distributions, with few differences between biopsies taken from upper, middle and lower lobes (Figure 1F, Figure E1B). The most striking variation was for submucosal gland cells (serous and mucous cells). Their detection in 3 out of 3 nasal biopsies, 3 out of 9 tracheal biopsies and 0 out of 9 intermediate biopsies (Figure E1B) suggests a larger density of glands in the nose, and a progressive decline in smaller airways, as previously described (17–20). Comparison between nasal and distal brushing samples also showed a clear enrichment of secretory cells in nasal samples, and an enrichment of multiciliated cells in distal samples (Figure 1F, Figure E1B). In order to characterize qualitative differences between nasal and tracheobronchial compartments, we assessed the correlations in average gene expression between each epithelial cell type. We found stronger correlations (>0.9) between cells belonging to the same cell type, in a donor-independent manner, than between cells belonging to distinct cell types (Figure 2A), confirming that cell type identity was well conserved across samples (Figure E8). This analysis also revealed nasal-specific and tracheobronchial-specific sub-clusters for suprabasal, secretory and multiciliated cells (Figures 1C, 1D, 2A, Figure E8), characterized by differentially expressed genes. We found a higher number of overlapping genes between suprabasal, secretory and multiciliated cell types from the nasal epithelium than between the same cell types from the tracheobronchial epithelium (Figure 2B and 2C). Among the top 22 genes shared by all 3 cell types in nasal samples were *SIX3*, *PAX7* and *FOXG1* (Figure E9), which have well-reported roles in the eye, neural and/or neural crest-derived development (21–23).

We then compared gene expression between secretory cells from nasal or tracheobronchial origins (Figure 2D). A total of 543 genes were found up-regulated in the nose with a FC>1.5, logCPM >1.5 and p-value <0.05, whereas 499 genes were found up-regulated in the trachea/bronchi with identical thresholds (Table E3). We noticed an enriched expression of *SCGB1A1*, *SCGB3A1*, *KLK11* and *SERPINF1* in the bronchi, and of *LYNX1*, *S100A4*, *CEACAM5*, *LYPD2*, *PI3* and *MUC4* in the nose (Figure 2D, Figure E9). Immunostainings confirmed the nasal expression of *PI3* and *MUC4* on independent brushing and biopsies (Figure 2F). Thirty seven additional transcripts were confirmed, based on a comparison with the Protein Atlas database (24) (Table E4). First insights about distinctive functional properties were obtained after a gene set enrichment analysis. In bronchi, functional terms related to defense and innate immune response to aggression were found. In nasal epithelium, enrichment in terms related to differentiation and motility supports the existence of a higher cellular turnover (Figure 2E, Table E5). Several regulatory units were associated with secretory cells of the nasal epithelium, such as *MESP1*, reported as a regulator of somitic mesoderm epithelialization (25). *AHR*, the aryl hydrocarbon receptor, which contributes to adaptive and innate responses by inducing the expression of several xenobiotic-metabolizing enzymes, and *STAT1*, a transcription factor that acts downstream to the interferon pathway, were both enriched in the nasal tissue. *FOXA3* regulatory unit, which promotes goblet metaplasia in mouse and induces *MUC5AC* and *SPDEF* expression (26, 27), was enriched in tracheobronchial samples.

Intriguingly, dissociated nasal cells appeared larger. There was a proximo-distal gradient of cell size, with the largest average size in the nose (12.56 µm ± 0.71) and the smallest size in the distal airways (8.77 µm ± 0.71) (Figure E2C and E2D). This difference correlated with the number of detected genes (Figure E2A and E2B).

### Identification of rare epithelial cells along the human airways

We identified 13 brush/tuft cells according to their high expression of *LRMP* and *RGS13* (12, 28, 29) (Figure 3A-3C). We also noticed in these cells a specific activity of *HOXC5*, *HMX2*, and *ANXA4* regulatory units (Figure E10A).

A cluster of 29 pulmonary neuroendocrine cells (PNECs) (Figure 3A) was found, mostly in tracheal and intermediate biopsies (Figure 3B). PNECs expressed the neurotransmitter-associated genes *PCSK1N*, *SCGN* and *NEB* (Figure 3C) and we identified *HOXB1*, *ASCL1*, and *FOXA2* as PNEC-specific regulatory units (Figure E10A).

A cluster of 117 ionocytes was also identified (Figure 3A), mostly in nasal and distal brushings (Figure 3B), indicative of their more luminal position. Ionocytes were characterized by markers such as *ASCL3* and *CFTR* (29) (Figure 3C), and we identified ASCL3, *FOXI1* and *DMRT2* as ionocyte-specific regulatory units (Figure E10A).

A cluster of 63 cells, labelled as “undefined rare” cells, was sampled evenly across all macro-anatomical locations (Figure 3A and 3B). Relative to the other rare populations, these cells expressed more specifically *NREP*, *STMN1* and *MDK* (Figure 3C) but shared the expression of *HEPACAM2*, *HES6*, *AZGP1*, *CRYM* and *LRMP* with ionocytes, brush cells and PNECs. When we searched for a correlation with the other epithelial cell types, we found a high correlation with ionocytes (>0.85), PNECs and brush cells (>0.80), and also with basal and suprabasal populations (>0.85) (Figure E10B). This profile suggests a precursor role for these cells.

A last group of rare cells, already described in primary cultures (12) and in asthmatic patients (30) named multiciliating-goblet cells, corresponded to cells expressing both goblet and multiciliated cell markers. In our dataset, around 60 cells were found positive for both *FOXJ1* and *MUC5AC*. They were equally distributed between the secretory and the multiciliated cell clusters. We have used SoupX to remove gene counts that may emerge from cell-free mRNA contamination, thus avoiding interference with the quantification of multiciliating-goblet cells (Figure 3E). We confirmed the presence of these cells using RNAscope *in situ* RNA hybridization (Figure 3F). When these cells were superimposed in a PAGA representation of tracheobronchial cell lineages, we noticed that they were located close to multiciliated cells, while in a similar representation of nasal cell lineages, they were located between secretory and deuterosomal cells, nearer to these latter (Figure E10B). This result supports our previous description of goblet cells as precursors of multiciliated cells in homeostatic and healthy epithelium and additionally suggests that transition through this stage may have slightly distinct dynamics between nasal and tracheobronchial epithelia (12).

### Cell proliferation within homeostatic airways

Before batch correction, we identified a cluster of cycling cells, defined by the expression of *MKI67*, *TOP2A*, *CDC20* (Figure 4A). After batch correction, these cells spread between the basal and suprabasal clusters (Figure 4B). A cell cycle analysis of all cell types identified 2 clusters with positive cell cycle scores. One corresponds to cycling cells (*MKI67*-positive) and the other, to deuterosomal cells (*MKI67*-negative) (Figure 4C), in agreement with Ruiz Garcia *et al.* (12). Figure 4D shows UMAP graphs for the subgroup of cells that belonged to the *bona fide* cycling cluster with a superimposition of the cell cycle scores for G1, S and G2/M phases, which delineates each phase of the cell cycle inside the circular embedding (Figure 4D). We noticed that the marker genes of this cycling population largely overlap with those of suprabasal cells (Figure 4E), suggesting that in the homeostatic and healthy epithelium, suprabasal cells may be the main proliferating population in the epithelium. Labelling of bronchial epithelium sections with *MKI67* antibody confirmed the presence of *MKI67*+/*KRT5*+ cells that were located in a para/suprabasal position (Figure 4F). Cycling cells were distributed all across the 35 samples, although with a highly variable distribution, which was reminiscent to the expression profile of *KRT13* in suprabasal cells (Figure 4G and 4H, Figure E11A). These *KRT13*-high samples also displayed the highest cycling cell proportion (more than 20% of cycling cells, Figure 4G). *In situ* RNA hybridization in nasal epithelium sections confirmed an association of *MKI67* RNA with cells expressing *KRT13* (Figure E11B). This association between *KRT13* expression and proliferation, together with the variability of detection of these cells, is highly reminiscent of the previous description of hillocks in mouse airway epithelium (29). We indeed confirmed the presence of *KRT13*+ cell clusters in nasal epithelium, with patterns very similar to those previously found in mouse (Figure 4I).

**Figure 4.**
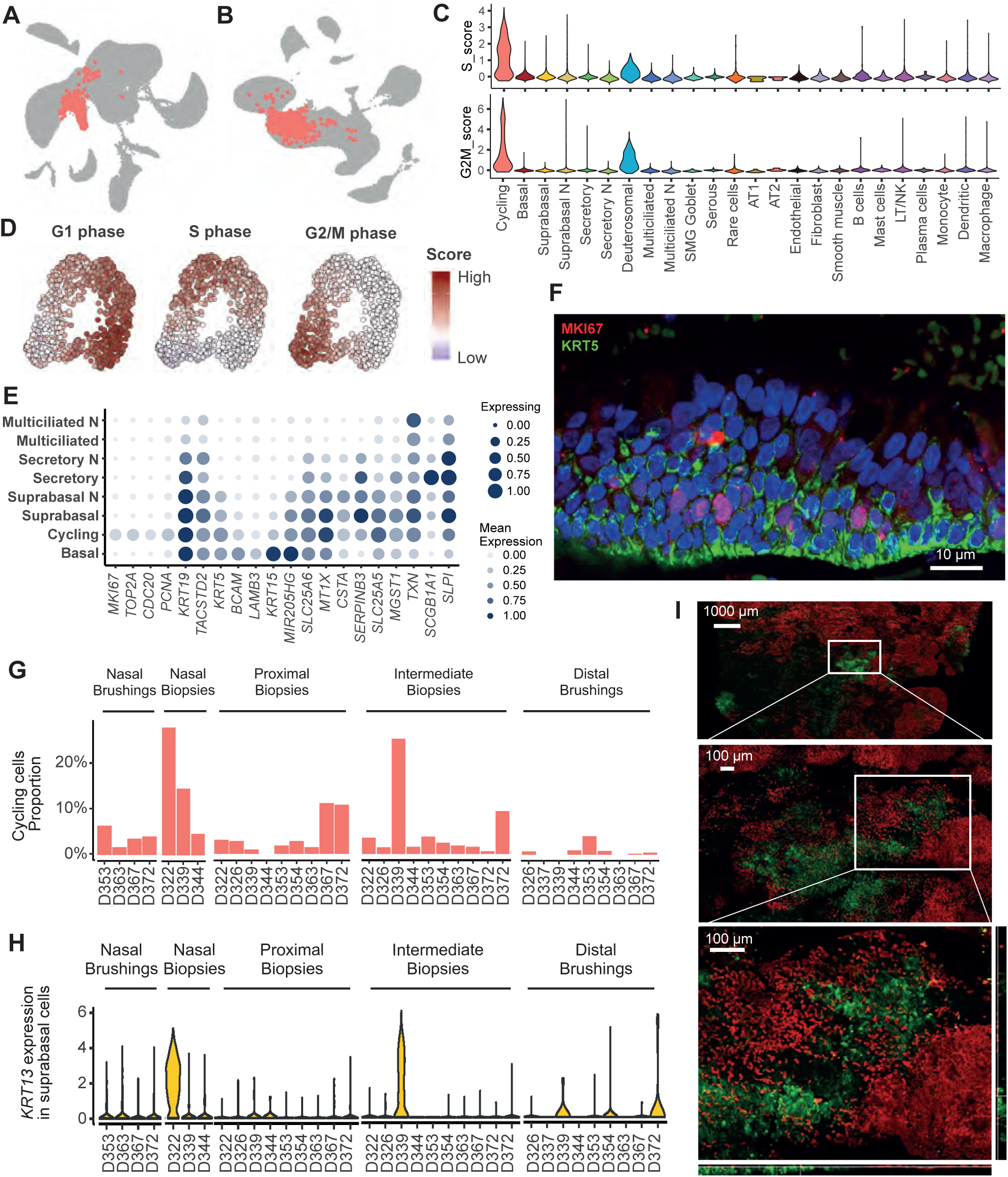
Characterization of cycling cells and KRT13 expression in the healthy airway epithelium. **(A-B)** Highlights of cycling basal cells in global UMAP representations without **(A)** or with **(B)** batch correction of the embedding. **(C)** Violin plot of the cell-cycle phase score in all cell types detected in the whole dataset. **(D)** Focused UMAP visualizations on the subset of cycling cells, colored by cell cycle phase scores at G1, S, G2/M stages. **(E)** Dot Plot of marker gene expression in cycling, basal and suprabasal cells. **(F)** Immunostaining for MKI67 and KRT5 in a bronchial biopsy section. **(G)** Barplot of the percentage of cycling cells per sample. **(H)** Violin plots of the expression of *KRT13* in suprabasal cells. **(I)** Immunostainings for KRT13 (green) and acetylated tubulin (red) in nasal turbinate whole mount (top view).

## Discussion

We have established a reference single-cell atlas of normal human airways after analyzing 35 fresh tissue samples collected by bronchoscopy in 10 healthy volunteers, resulting in a large-scale gene expression profiling that also integrated spatial information of each sample. This approach was well adapted to collect samples from the nose to the mid airways but excluded the bronchiolar compartment and the parenchyma, for which alternative experimental approaches have already been proposed (30, 31). Establishing a comprehensive airway atlas will result from the combination of our atlas with these other datasets. That saying, our approach provides a unique opportunity to build a single-cell gene expression resource based on well-characterized healthy volunteers which are rarely accessible in most large scale studies. The use of bronchoscopy, a minimally invasive approach to the airways, also creates a real opportunity to transfer rapidly novel information generated in the context of the Human Cell Atlas project to new clinical practices.

In our workflow, a critical analytical step led to a robust cell type annotation of 35 single-cell RNA-sequencing experiments. Integration was performed sequentially, after quantification of individual samples, their merging and batch correction. The quality of our sampling and analysis resulted in non-significant donor-related effects and in unprecedented high epithelial cell proportions, both important quality criteria which had not been systematically reached by the other lung atlas reports, making our resource the most reliable one so far. Our conclusions were all based on observations which were made on several donors and independently confirmed. For instance, our description of rare epithelial cell types was based on information collected from all 10 donors.

Profiling of identical cell types across many sites of the airway tree has allowed us to quantify the frequencies of epithelial, submucosal-gland, immune and stromal cells, and has revealed an influence of the mode of sampling. However, this did not prevent us from defining stable core cell type signatures for each epithelial, stromal and immune cell types, irrespective of their anatomical location. In contrast, important variations of gene expression were found when comparing the same populations of suprabasal, secretory and multiciliated cells from the surface epithelium between nasal and tracheobronchial compartments. Some variations, which did not reach significance, were also found in basal cells. These results fit well with previous works reporting dozens of differentially expressed transcripts between nasal and bronchial brushings (32, 33). We also tested whether additional gradients of expression could be detected, for instance between upper and lower lobes, or between anterior or posterior part of the trachea. While we noticed some trends, the observed differences, such as an increased number of T cells, B and dendritic cells in lower lobes, need additional measurements to be validated. Future functional analysis will be important to assess the impact of these variations on the biology of each cell type. Our gene set enrichment and inferred regulatory unit analysis suggest that nasal cells undergo higher epithelial regeneration, and have function in xenobiotic metabolism as well as in interferon signaling. Interestingly, *SIX3*, *PAX6-7*, and *OTX1/2*, which we found to be specific of the nasal epithelium, are all associated with gene ontology terms such as “pattern specification process”, and “axis specification” and have well-described functions during embryonic patterning of the head (22, 34–36). Expression of *Six3* in murine ependymocytes, which are radial glia-derived multiciliated cells, is necessary for the maturation of these cells during postnatal stages of brain development (34). Hence, nasal-specific expression of developmental patterning genes might be the consequence of head vs. trunk differential developmental origins and may not necessarily confer specific functions to nasal epithelial cells. The underlying mechanisms that confer a persistence in the expression of these developmental hallmarks remain to be elucidated.

Our focus on secretory cells demonstrates that nasal ones contain few *SCGB1A1^+^*- and *SCGB3A1^+^*-cells. Despite this low secretoglobin content, they display the core gene signature of secretory cells, suggesting that secretoglobins may not be sufficient marker genes to identify all secretory/club cells. These differences are important to consider when using nasal samplings as a proxy to assess bronchial status.

Our atlas also provides a comprehensive description of novel cell types, such as the multiciliating-goblet cells and the undefined rare cells, which, given their gene expression profile, may be precursors for the ionocytes, PNECs and brush cells. This is, to our knowledge, the first scRNAseq identification of human PNECs and brush cells with distinct gene signatures. We also identified *KRT13^+^*-cells with a detection frequency and organization that are highly reminiscent of the mouse airway hillocks (29), providing the first identification in human airways of hillock structures, and further work is required to investigate their functions.

Altogether, this atlas improves the cellular stratification of gene expression profiles in healthy human airway epithelium. It now makes possible an extensive exploration of the various situations involved in homeostasis and regeneration of normal and pathological airways.

## Acknowledgements

We are grateful to the UCAGenomiX platform for fruitful discussions and technical help on single-cell RNA sequencing, to Frederic Brau, Sophie Abelanet and Julie Cazareth from the IPMC imaging platform, for fruitful discussions and technical help on imaging and to Jennifer Griffonnet for her invaluable help in building an experimental protocol that respects ethical rules, in collaboration with the French health authorities (Agence Nationale de Sécurité du Médicament et des produits de santé and Comité de Protection des Personnes).

## Online Materials and Methods

### Ethics statement

The study was approved by the Comité de Protection des Personnes Sud Est IV (approval number: 17/081) and informed written consent was obtained from all participants involved. All experiments were performed during 8 months, in accordance with relevant guidelines and French and European regulations. No deviations were made from our approved protocol named 3Asc (An Atlas of Airways at a single cell level - ClinicalTrials.gov identifier: NCT03437122).

### Human samples

Human samples were collected from healthy adult volunteers during bronchoscopy under local anaesthesia. All procedures were administered by the same pulmonologist at Nice university hospital, France. The process, the location and type of specimens (brushing or biopsy) were compatible with future use in daily clinical practice. Samples were taken at distinct levels of the respiratory tract: nose (lower turbinate), trachea (carina), intermediate bronchi (5^th^-6^th^ divisions) and distal (9^th^ to 12^th^ divisions). Intermediate and distal samples were taken to obtain, with all subjects included, the broadest mapping in terms of upper, middle and lower pulmonary segments. The description of each sample can be found in Table E1.

### Sample dissociation

All sample dissociation protocols are available on protocols.io : brushings (protocol qubdwsn), biopsies (protocol x3efqje).

#### Dissociation of brushings (protocols.io qubdwsn)

The brush was soaked in a 5 mL Eppendorf containing 1 mL of dissociation buffer which was composed of HypoThermosol® (BioLife Solutions) 10 mg/mL protease from Bacillus Licheniformis (Sigma-Aldrich, reference P5380) and 0.5 mM EDTA. The tube was shaken vigorously and centrifuged for 2 min at 150 g. The brush was removed, cells pipetted up and down 5 times and then incubated cells on ice for 30 min, with gentle trituration with 21G needles 5 times every 5 min. Protease was inactivated by adding 200 μL of inactivation buffer (HBSS/2% BSA). Cells were centrifuged (400g for 5 min at 4°C). Supernatant was discarded leaving 10 μL of residual liquid on the pellet. All subsequent centrifugation and supernatant removal steps have been performed following the same procedure. Cells were resuspended in 200 µL of wash buffer (HBSS + 1% BSA). Cells were observed under an inverted microscope and red blood cells (RBC) content was evaluated with a Countess FL II automated cell counter (Thermo Fisher Scientific), after addition of Hoechst 33342 to an aliquot of the cell suspension to discriminate nucleated cells from non-nucleated cells. RBC lysis was performed if RBC content was higher than 50%. Prior to RBC lysis, cells were centrifuged and resuspended in 100 µL PBS. 900 µL (9 volumes) of Ammonium Chloride 0.8% (StemCells technologies,07800) were added to 100 µL of cell suspension. Following a 5 min incubation on ice, 400 µL of inactivation were added and cells were centrifuged. Cells were resuspended in 1000 μL of wash buffer and passed centrifuged again. If no RBC lysis was performed, this was the final wash. If RBC lysis was performed, one additional wash step was performed. Before last centrifugation, cells were passed through 40 µm porosity Flowmi™ Cell Strainer (Bel-Art). Cells were resuspended in 30 μL of wash buffer. Cell counts and viability were performed with Countess™ automated cell counter (Thermo Fisher Scientific). For the cell capture by the 10X genomics device, the cell concentration was adjusted to 500 cells/µl in HBSS aiming to capture 5000 cells. All steps were performed on ice.

#### Dissociation of bronchial biopsy (protocols.io x3efqje)

The biopsy was soaked in 1 mL dissociation buffer which was composed of DPBS, 10 mg/mL protease from Bacillus Licheniformis (Sigma-Aldrich, reference P5380) and 0.5 mM EDTA. After 1 h, the biopsy was finely minced with a scalpel, and returned to dissociation buffer. From this point, the dissociation procedure is the same as the one described in the “dissociation of brushings” section, with an incubation time increased to 1h. For the cell capture by the 10X genomics device, the cell concentration was adjusted to 500 cells/µl in HBSS aiming to capture 5000 cells. All steps were performed on ice.

#### Cytospins from brushings

Cells dissociated from brushings were cytocentrifuged at 72 g for 10 min onto SuperFrostTM Plus slides using a Shandon CytospinTM 4 cytocentrifuge. CytospinTM slides were fixed for 10 min in 4% paraformaldehyde at room temperature for further immunostaining.

### Tissue handling for immunostaining and in situ RNA hybridization

#### Processing of nasal turbinates

Inferior turbinates were resected from patients who underwent surgical intervention for nasal obstruction or septoplasty (kindly provided by Professor Castillo, Pasteur Hospital, Nice, France). The use of human tissues was authorized by the bioethical law 94–654 of the French Public Health Code after written consent from the patients. After surgery, nasal inferior turbinates were immediately immersed in Ca2+/Mg2+-free HBSS supplemented with 25 mM HEPES, 200 U/ml penicillin, 200 μg/ml streptomycin, 50 μg/ml gentamicin sulfate and 2.5 μg/ml amphotericin B (all reagents from Gibco). After repeated washes with ice-cold supplemented HBSS, tissues were processed depending on the assay.

#### Whole mounts of nasal turbinate epithelium

The outer layer (approximatively 1.5-mm thick) of nasal turbinates was resected with the help of a scalpel blade allowing the recovery of the epithelium that covers the turbinates. Nasal epithelium was fixed in PFA 4% for 1 hour at room temperature then overnight at 4°. After two washes in PBS, the epithelium was permeabilized with 0.5% Triton X-100 in PBS, blocked in 0.3% BSA for 30 min. Primary antibodies were incubated for 24 hours at room temperature, washed in 0.3% Triton X-100 in PBS, incubated with appropriate secondary antibodies diluted in blocking buffer for 4 hours at room temperature, washed in 0.3% Triton X-100 in PBS, all the steps were performed in a shaker. The epithelium was then mounted between a slide and cover-slip using imaging spacers. Imaging of the samples was performed in a Confocal LSM780 Zeiss.

#### Cryostat section of nasal turbinate epithelium

Nasal turbinates were fixed in paraformaldehyde 4% at 4°C overnight then extensively rinsed with phosphate-buffered saline (PBS). Fixed tissues where then prepared for cryo-embedding for cryostat sectioning. Tissue was embedded in optimal cutting temperature (OCT) medium (Thermo Fisher Scientific) at room temperature and then frozen by contact with liquid nitrogen. 10 µm-thick frozen tissue sections were obtained with a cryostat Leica CM3050S on Superfrost Plus® Gold slide (Thermo Scientific). Sections were kept at -80°C with desiccant for few weeks until use for RNAscope protocol.

### RNA *in situ* hybridization with RNAscope

#### Pretreatment Protocol

For cryostat tissue sections, the manufacturer’s protocol for fixed frozen tissues described in user manual RNAscope® Multiplex Fluorescent Reagent Kit v2 Assay (Cat. No. 323100, Advanced Cell Diagnostics, lnc., USA) was followed. To avoid tissue section detachment from slides, the target retrieval step was replaced by an increased protease III incubation time to 45 min. For cytospin samples, the cell pretreatment described in ACD technical note MK-50 010 was followed. As red blood cell lysis has been performed during cell dissociation, hydrogen peroxide treatment step was skipped in further pretreatment. Protease III was incubated 30 min without dilution as cytospin cells are fixed with paraformaldehyde to follow the same pretreatment condition described by ACD for fixed frozen tissue (Cat. No. 323100, Advanced Cell Diagnostics, lnc., USA).

#### RNAscope Assay

After pretreatment, for both sections and cytospin samples, we followed manufacturer’s instructions for RNAscope® 4-plex Ancillary Kit for Multiplex Fluorescent Reagent kit v2. Briefly, 20 double Z probe pairs specifically targeting the region coding for each targeted genes were designed and synthesized by ACD. ACD probes used were: FOXJ1-C1 (430921), SCGB1A1-C4 (469971-C4), MUC5AC-C2 (312891-C2), KRT13-C1 (528111), KRT5-O1-C2 (547901-C2), MKI67-C3 (591771-C3). Hybridization signals were detected by Opal probes 520, 570 and 650 (Cat. No. FP1487001KT, FP1488001KT and FP1496001KT, Perkin Elmer) at 1:1500 dilution. At last, sections on glass slides were counterstained with DAPI for 30 sec and mounted in Prolong^TM^ Gold antifade reagent with DAPI (Cat. No. P36931, Life technologies). The images were captured by a Zeiss LSM780 confocal microscope.

### Immunostaining of paraffin sections

Sections were deparaffinized, an antigen retrieval treatment was performed using citrate buffer at pH6. Sections and cytospins were permeabilized with 0.5% Triton X-100 in PBS for 10 min, a following blocking treatment was performed with 3% BSA in PBS for 30 min. The incubation with primary antibodies was carried out at 4°C overnight. Incubation with secondary antibodies was carried out during 1h at room temperature. Nuclei were stained with 4,6-diamidino-2-phenylindole (DAPI). Primary and secondary antibodies information represented in supplementary table E6. When necessary, KRT5 antibody was directly coupled to CF488 with the Mix-n-Stain kit (Sigma-Aldrich) according to the manufacturer’s instruction. Coupled primary antibody was applied for 2 hours at room temperature after secondary antibodies had been extensively washed and after a 30 min blocking stage in 3% normal rabbit serum in PBS. Imaging of the samples was performed using a Confocal FV-10 from Olympus.

**Table E6.**
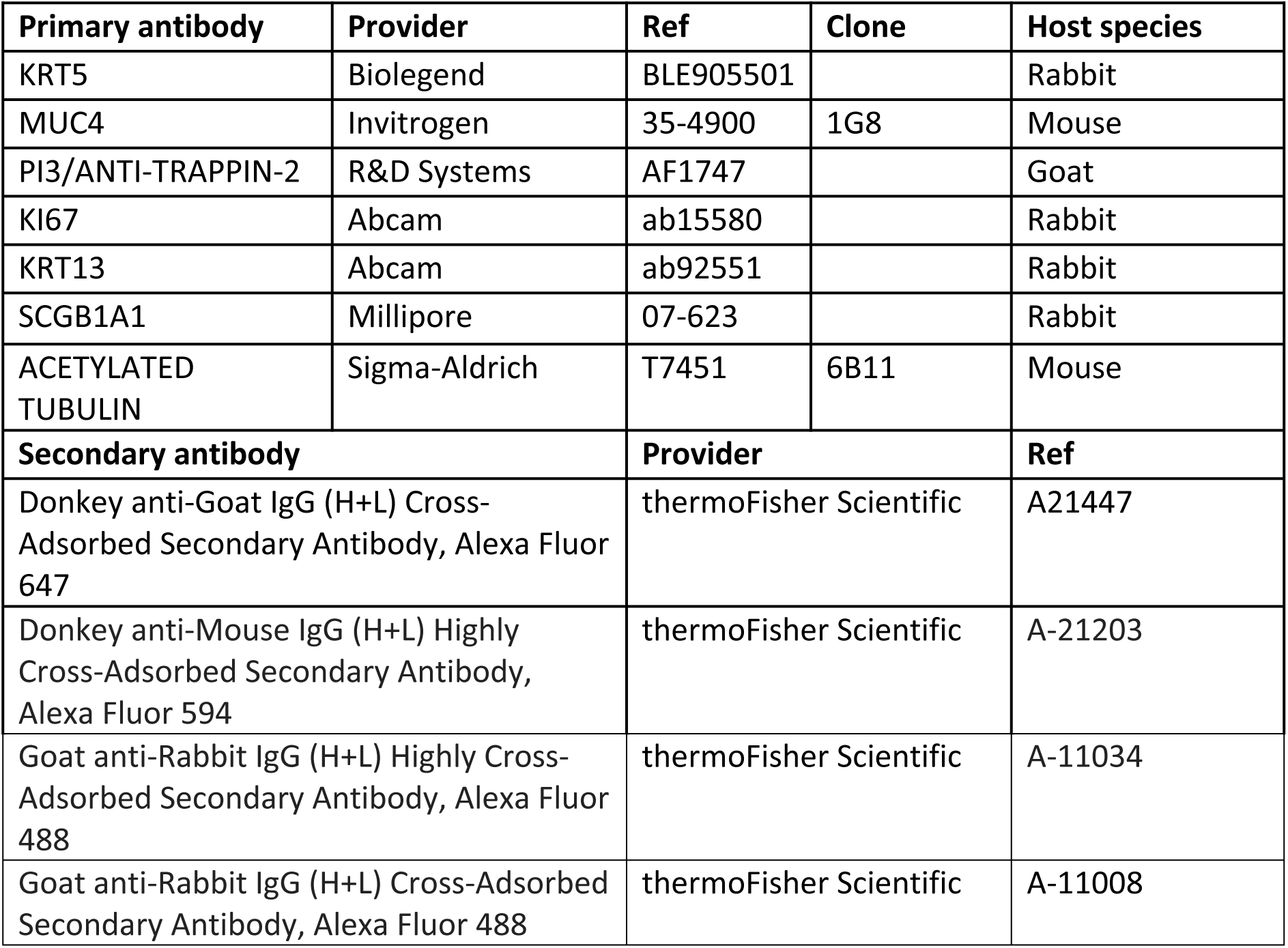
Antibody used for immunostainings.

### Chromium 10X Genomics library and sequencing

We followed the manufacturer’s protocol (Chromium™ Single Cell 3’ Reagent Kit, v2 Chemistry) to obtain single cell 3’ libraries for Illumina sequencing. Libraries were sequenced with a NextSeq 500/550 High Output v2 kit (75 cycles) that allows up to 91 cycles of paired-end sequencing: Read 1 had a length of 26 bases that included the cell barcode and the UMI; Read 2 had a length of 57 bases that contained the cDNA insert; Index reads for sample index of 8 bases. Cell Ranger Single-Cell Software Suite v2.3.0 was used to perform sample demultiplexing, barcode processing and single-cell 3′ gene counting using standards default parameters and human build hg19. All single-cell datasets that we generated, and the corresponding quality metrics are displayed in Table E1.

### Primary data analysis

Initially, each dataset was roughly analysed using Seurat (v3)^[1]^ to determine the best analysis workflow needed for the merged dataset. Permissive filtering was done on low-quality cells followed by median normalization, identification of highly variable genes and Louvain clustering. Marker genes of cell-clusters were identified using Wilcoxon’s rank test and shared top genes across datasets resulted in a common and robust cell type annotation. This initial annotation was later used to create a reference for the precise annotation of the merged dataset (Figure E8).

### Data quality control done on individual datasets

#### Cell and gene filtering

Each sample processing being slightly different from others (sample size, presence of blood, dissociation times), sample-specific quality metrics vary slightly between samples. To take into account this variability, each sample was pre-processed individually. Cells were excluded based on three criteria: high number of Unique Molecular Identifier (UMIs) per cell (max +3 Median Absolute Deviation, MAD), low number of detected genes per cell (min 500 genes) and high percentage of mitochondrial genes (max +3 MAD). Mitochondrial and ribosomal genes (gene symbols starting with RPS/RPL) were excluded from the count matrices.

#### Doublet removal

We used DoubletDetection for unbiased identification of doublets (technical error) in our datasets (https://github.com/JonathanShor/DoubletDetection). We kept the default parameters values and removed the predicted doublets. Further doublet removal was done on the merged dataset (without data integration), to remove clusters with a high proportion of predicted doublet (over 50 %).

#### Ambient mRNA correction

Dissociation of complex tissue, such as brushing and biopsies, results in a certain proportion of cell lysis. It results in the presence of ambient mRNA that spreads across all droplets of a single experiment. This gene expression background is highly dependent of the cell-type composition which might lead to misleading analysis. We used soupX (https://github.com/constantAmateur/SoupX) for background correction and compared analysis results between corrected and uncorrected dataset (corrected dataset being very sparse). Background corrected data were mainly used for visualization purposes.

### Data integration

#### Normalization

Size factors were calculated for the complete (merged) dataset using ‘ComputeSumFactor’ from the scran R package^[2]^. Cells were pre-clustered with the ‘quickCluster’ function, method ‘igraph’ and minimum and maximum cluster size of 100 and 3,000, respectively. Raw counts were then normalized and log-transformed with cell-specific size factors. A count of 1 was added to each value prior to log transformation.

#### Selection of highly variable genes

Highly variable genes (HVGs) were identified/calculated using the getHVGs function from the scran package, with default parameter values.

#### Batch Correction and Data Integration

Batch effects were removed using the ‘fastMNN’ function in the scran R package on 50 principal components computed from the HVGs only^[3]^. Correction was performed incrementally from the most homogeneous samples to the most heterogeneous (in QC and cell composition). Datasets were first corrected between sampling position and method (nasal biopsies, nasal brushings, tracheal biopsies, intermediate biopsies and distal brushings), merging sequentially from the sample containing the most cells to the samples containing the least. Positions were merged as follow: intermediate, tracheal, distal, nasal. The resulting batch-corrected principal component analysis was then used for further analysis steps. The compared analysis was performed between batch-corrected and uncorrected datasets.

### Dimensionality reduction and visualization

UMAPs were calculated using scanpy ^[4]^. For the complete dataset, the first 12 components of the batch-corrected PCA were used and considering the 100 nearest neighbours of each cell. For each data subset (immune, rare cells, stromal, and cycling) UMAPs were computed on uncorrected PCA based on subset specific HVGs.

### Data clustering and sub-clustering

We clustered cells using phenograph^[5]^ (available in scanpy) with two parameter settings (i: 12 PCs and 100 nearest neighbours) to tackle the imbalance in cell proportion (e.g. basal cells vs rare cells). The number of PCs used was estimated empirically on the PCA elbow plot, and by manual examination of top genes correlated with PCs. After a first annotation step, described below, a sub-clustering step was performed for each annotated cell type, to clean the boundaries between the distinct cell types (basal and suprabasal), but also in order to better identify small clusters such as rare cells or stromal cells. The number of PCs used for these sub-clustering steps varies from 3 to 8 with 20 nearest neighbours per cell.

### Markers identification and data annotation

Marker genes were identify using rank_genes_group function from scanpy using the Wilcoxon’s rank test. The robustness of those markers was assessed by reviewing the literature, and by the high correlation of phenograph clusters sharing similar markers genes. These clusters were then grouped and annotated as a unique cluster.

### Gene expression differential analysis

Differential analysis between specific clusters (secretory vs. secretory N for instance) was designed differently from the rank_gene_group function from scanpy to overcome the sample-specific gene expression background still present even after correction and to amplify the statistical power of the differential analysis. Pseudo-bulk samples were created from each cell-clusters by summing the raw counts of each gene in multiple single cells. Each bulk was designed to be composed of an equal number of cells (to get similar library size between bulks), and to contain randomly picked cells from a homogeneous mix of all the donor samples (to have a similar gene expression background between all bulks from the same cell type). Differential analysis was performed using glmFit function from the R package edgeR^[6]^.

### GSEA analysis

Gene set enrichment analysis was performed using the fgsea R package with the GO Biological Process gene sets (Broad Institute GSEA MSigDB).

### Cycling cell identification and cell cycle analysis

Cycling cells were identified in the batch-uncorrected analysis of the dataset as a single cluster, and this specific cell type annotation was reported in the batch-corrected dataset. Cell cycle scoring (S phase and G2M phase) was performed using the function score_genes_cell_cycle from scanpy tools and the associated cell cycle genes^[1]^. The G1 phase score was estimated as the opposite of both the S and G2M phase.

### Trajectory inference using PAGA

To compare the cell trajectories between nasal and tracheobronchial samples, two subsets of randomly picked cells from each nasal or tracheobronchial surface epithelial cell types (n = 500 cells per cell type) were used to infer their trajectories using the PAGA algorithm^[7]^ available in scanpy. The included epithelial cell types were cycling, basal, suprabasal, suprabasal N, secretory, secretory N, deuterosomal, multiciliated and multiciliated N cells. Cells were then projected on the corresponding force atlas embedding and multiciliated-goblet cells were highlighted on the resulting trajectory.

### Inference of transcriptional regulatory units

We inferred transcriptional regulatory units using the GRNboost2 algorithm implemented in the arboreto package (https://arboreto.readthedocs.io/en/latest/). Expression correlations between transcription factors and potential target genes were computed from a raw count data matrix where we set a maximum threshold of 5000 cells by cell types. We obtained 1222 modules composed of the 50 first top correlated genes with a confirmed transcription factor. We scored the activity of those modules in each cell of the complete dataset using the score_genes function from scanpy tools. Cell type-specific activity of each module was determined with a Wilcoxon’s rank test.

## Data availability

All data are currently submitted to the European Genome-phenome Archive (EGAS00001004082), and will then linked to the Data central Repository of the Human Cell Atlas, in order to ensure the openness of information. Data is also available through a dedicated web interface (https://www.genomique.eu/cellbrowser/HCA/)

## Supplemental table description

**Table E1. Sample information.** Description of each sample donor and anatomical location of origin, dissociation and quality control metrics of sequencing.

**Table E2. Number of cells per sample per cell type.** Description of the number of cells from each sample distributed across all identified cell types.

**Table E3. Nasal versus tracheobronchial differentially expressed genes for suprabasal, secretory and multiciliated cells.** Table of the differentially expressed genes between suprabasal and suprabasal N (a), secretory and secretory N (b) and multiciliated and multiciliated N (c). Logarithm based-2 of the fold change in gene expression between each group (LogFC). Average logarithm based-2 of the ‘Count Per Million’ of each gene (level of expression, mean logarithm of the sum of UMIs in synthetic bulks, cf. Methods). Likelihood ratio statistics applied during differential expression testing (LR).

**Table E4. Enriched gene sets associated with nasal and tracheobronchial secretory cells.** P-values and adjusted p-values are presented as well as the Enrichment Score (ES) and Normalized Enrichment Score (NES). The number of times a random gene set had a more extreme enrichment score value is described in the n_more_extreme column with the size of the gene set and the genes identified in the leading edge.

**Table E5. Validation in Protein Cell Atlas.** Thirty seven proteins coded by transcripts which were differentially expressed between nose and tracheobronchial cells were immunodetected in nasopharynx and bronchi by the Protein Atlas consortium. Quantification levels were extracted from the website (https://www.proteinatlas.org/about/download). Only the largest differential in both experiments are shown.

**Table E6. Inferred activity of regulatory units.** Top table of the regulatory unit activity per cell type (identified by Wilcoxon’s rank test) (a). Top 50 co-regulated genes composing a regulatory unit, ranked by correlation (b).

**Table E7. Cell type marker genes.** Top table of the differentially expressed genes (identified by Wilcoxon’s rank test in a one-cluster-vs-all design) for each cell type.

## Supplemental Figure Legends

**Figure E1.**
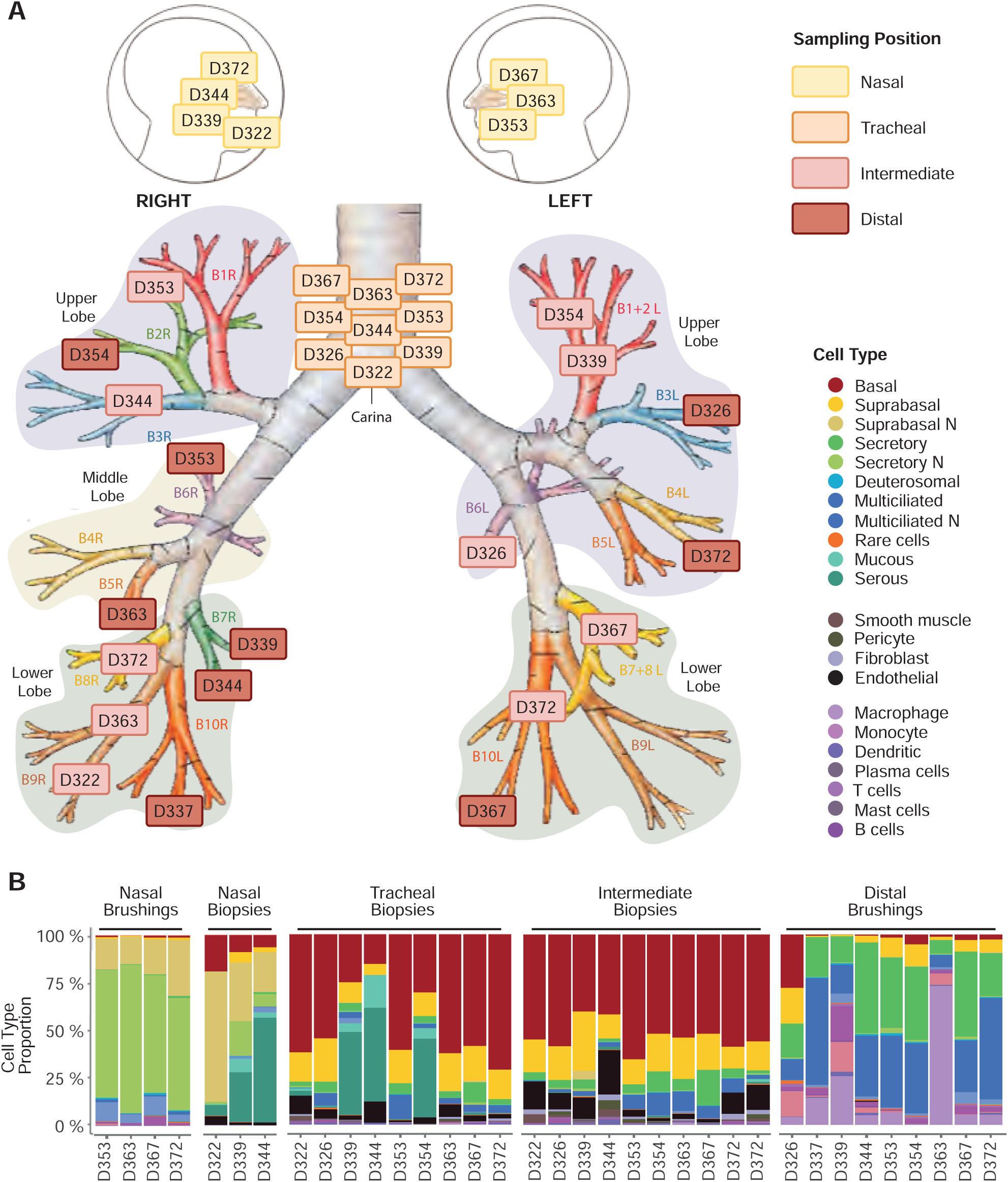
Sampling positions and cell type composition of the airways. (A) Schematic representation depicting the precise macro-anatomical location of each sample in the dataset. Numbers indicate donor identification number. (B) Barplot of the relative cell type composition of each sample, grouped by position and method of sampling.

**Figure E2.**
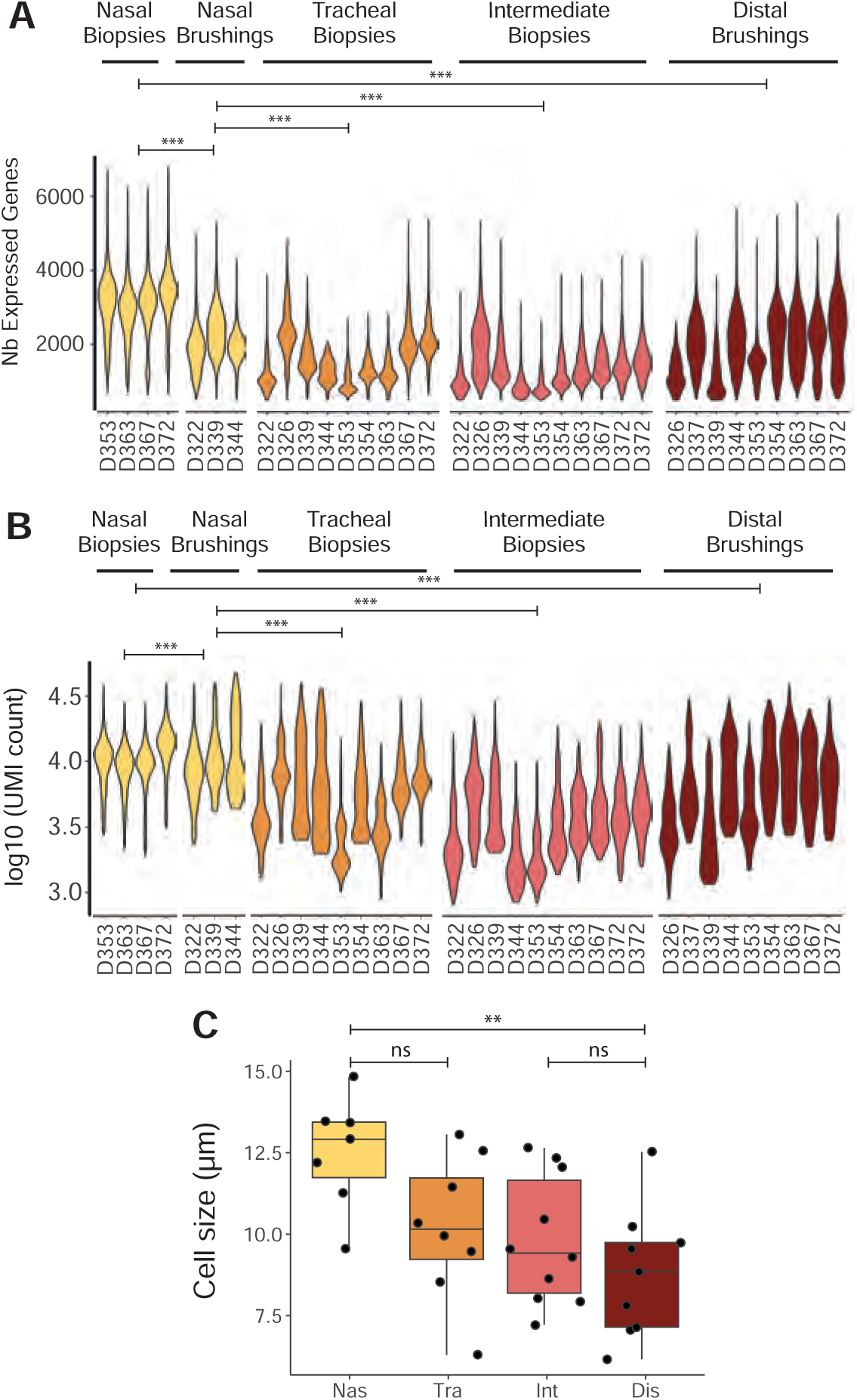
mRNA content and cell size of each sample. (A) Violin plot of the number of detected genes per sample. (Student t-test ***: pval < 0.001). (B) Violin plot of the number of UMI (log10 scale) per sample. (Student t-test ***: pval < 0.001). (C) Boxplot of average measured cell size per sample grouped by position (Wilcoxon test **: pval < 0.01, ns: non-significant).

**Figure E3.**
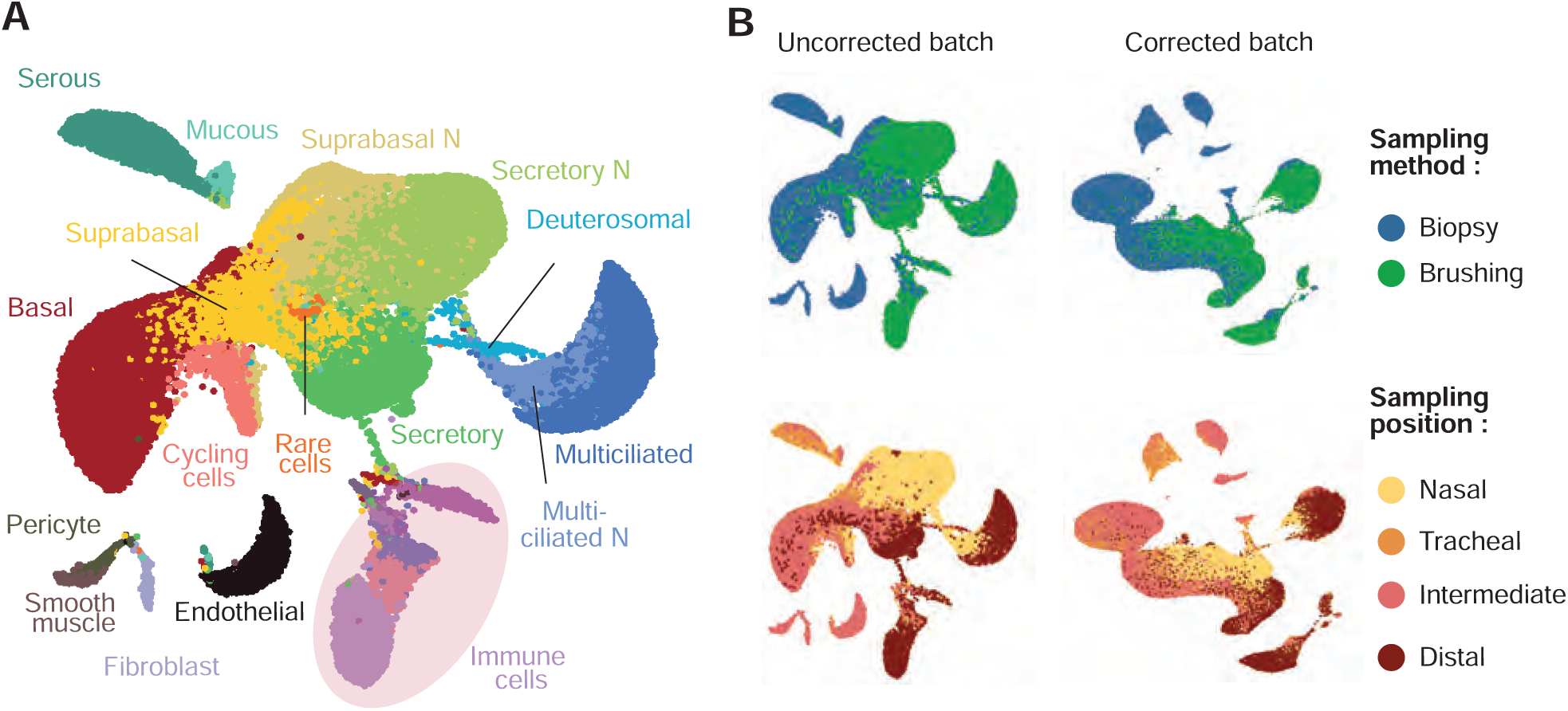
Dataset embedding. (A) UMAP visualization colored by cell types, without embedding. (B) UMAP visualization of batch corrected and non-batch corrected data, colored by cell type, sampling method or sampling position.

**Figure E4.**
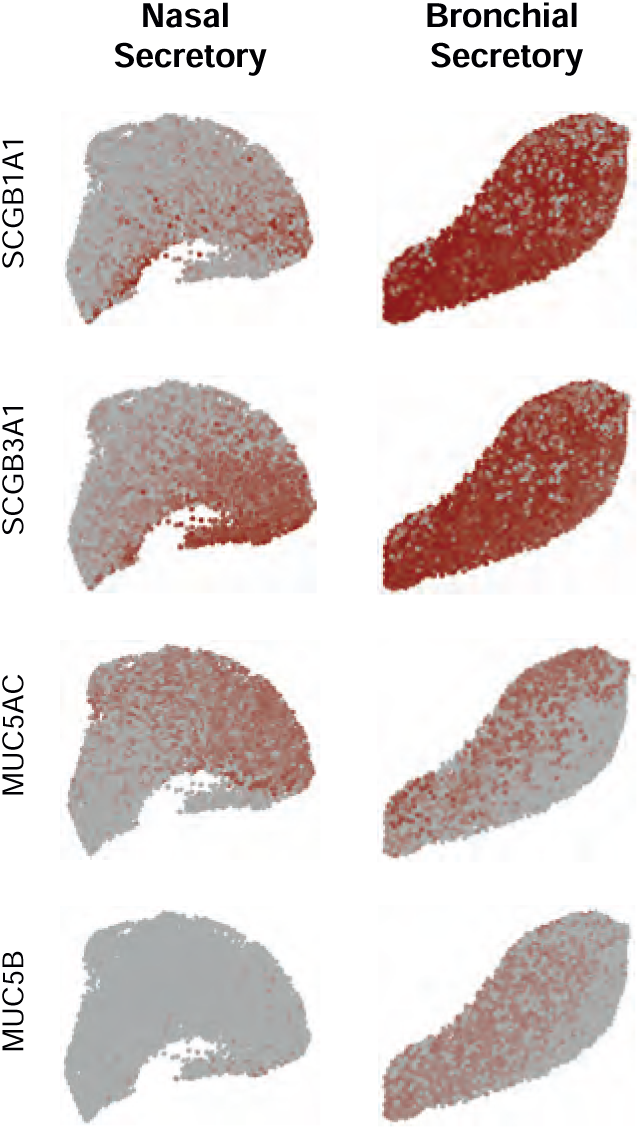
Secretory genes expression in secretory and secretory N cells. UMAP representation of secretory N (left) and secretory (right) cells for the selected genes.

**Figure E5.**
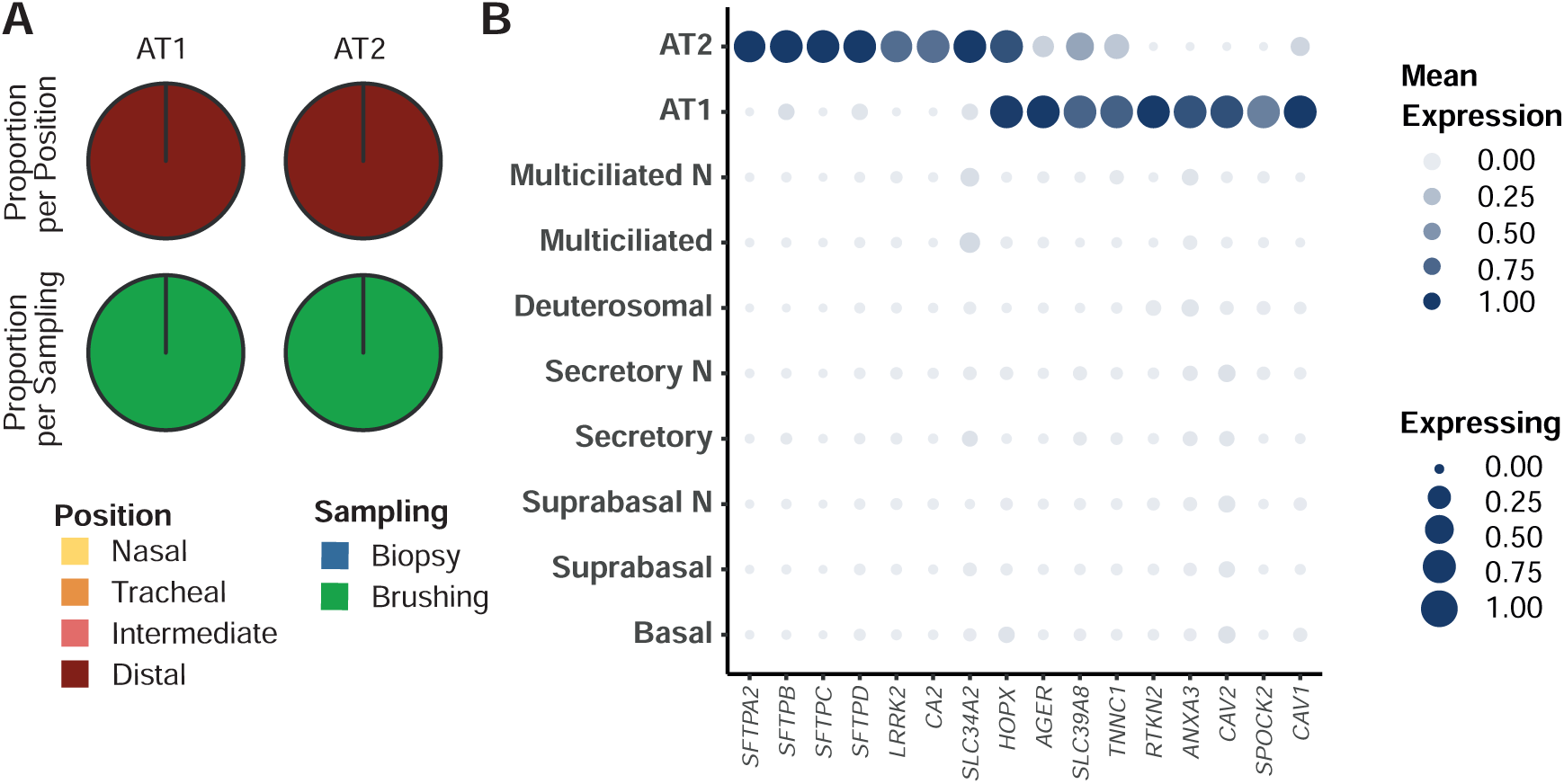
Pneumocyte distribution and characterization. (A) Pie chart of the anatomical region of origin for AT1 and AT2 cells (B) Dot plot of pneumocyte marker genes.

**Figure E6.**
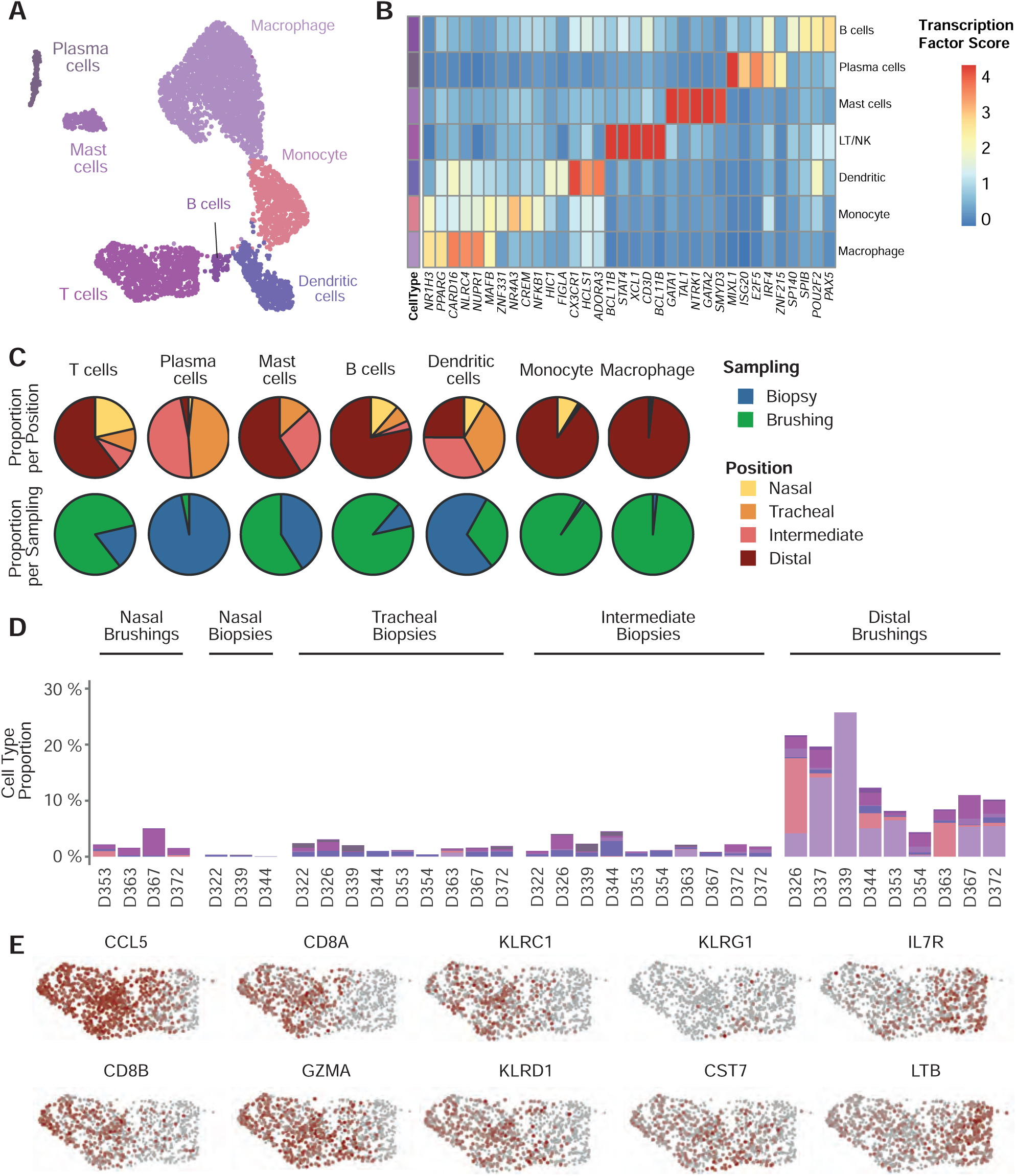
Immune cell distribution across the human airways. (A) UMAP visualization of the immune cell clusters. (B) Heatmap of cell type-specific regulatory unit activity score. (C) Pie chart of the anatomical region of origin for each immune cell type. (D) Barplot of the relative immune cell type composition of each sample, grouped by sampling position and method. (E) UMAP representation of T cells coloured by the expression of T cell subtypes marker genes.

**Figure E7.**
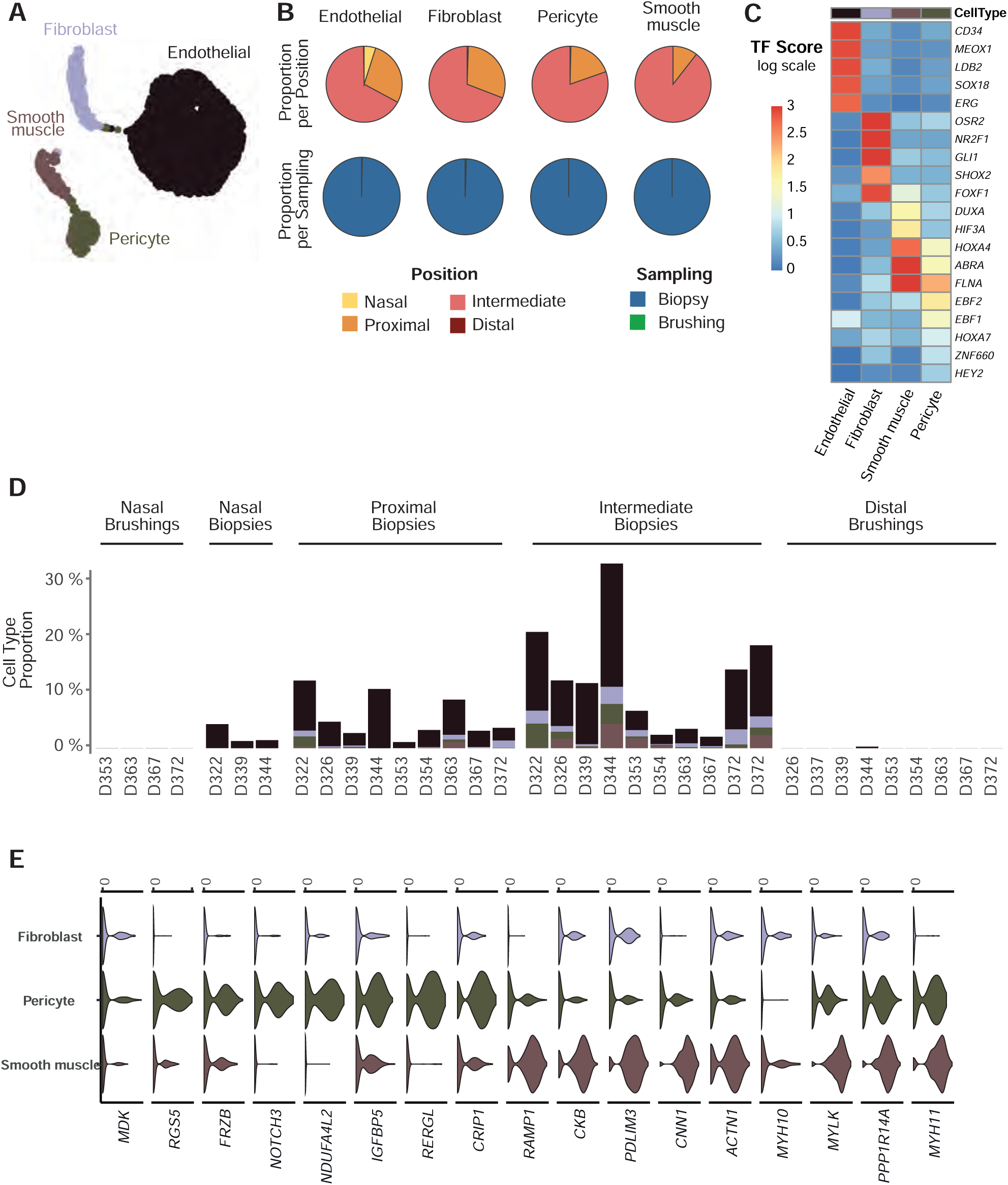
Mesenchymal cell composition across the human airways. (A) UMAP visualization of the mesenchymal cells cluster. (B) Pie chart of the anatomical region of origin for each mesenchymal cell type. (C) Heatmap of cell type-specific regulatory unit activity score. (D) Barplot of the relative mesenchymal cell type composition of each sample, grouped by sampling position and method. (E) Violin plot of marker genes associated with smooth muscles cells and pericytes.

**Figure E8.**
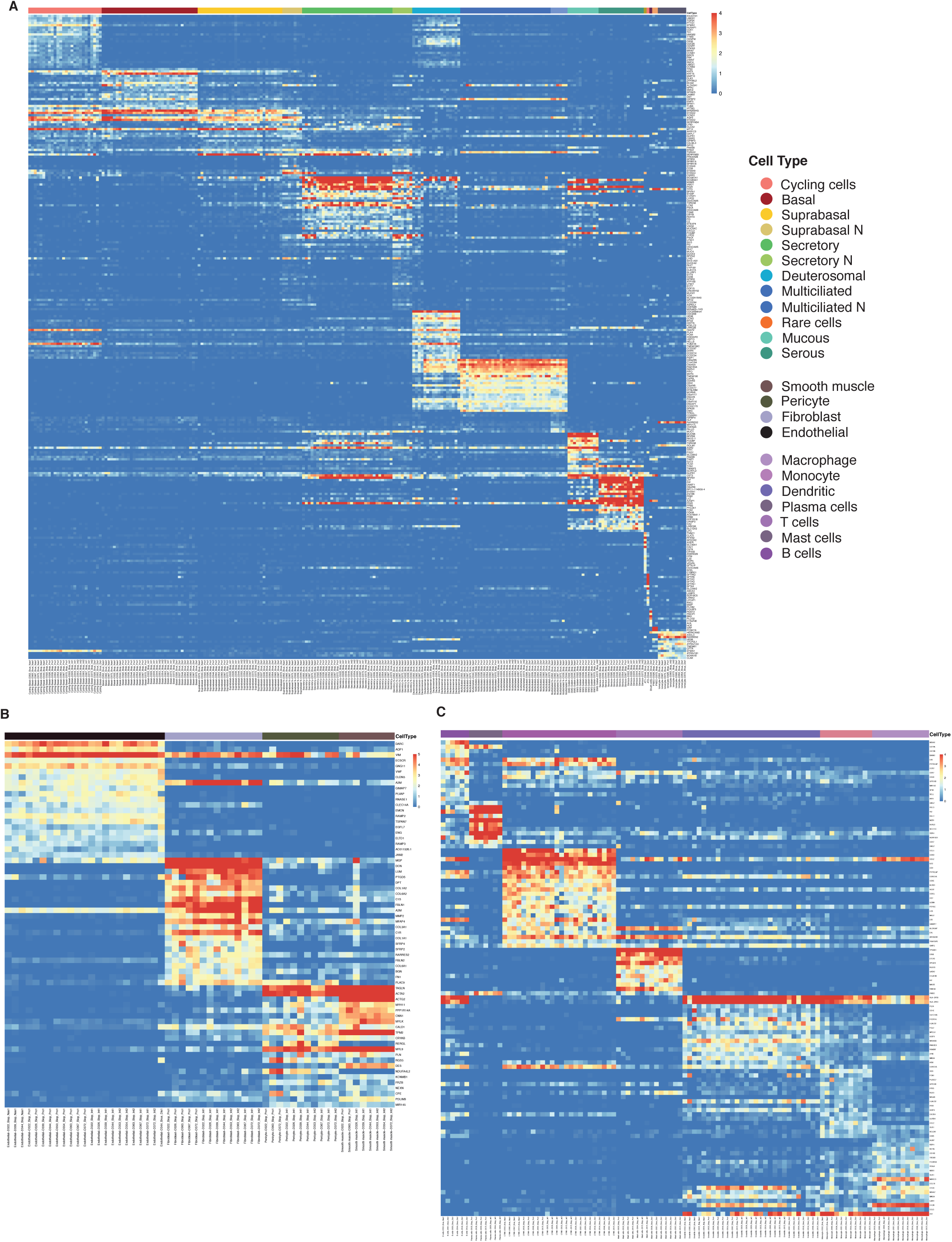
Robust heatmaps of cell type markers across all samples. (A) Heatmap of epithelial cell types. (B) Heatmap of stromal cell type. (C) Heatmap of immune cell types. Scaled by gene expression.

**Figure E9.**
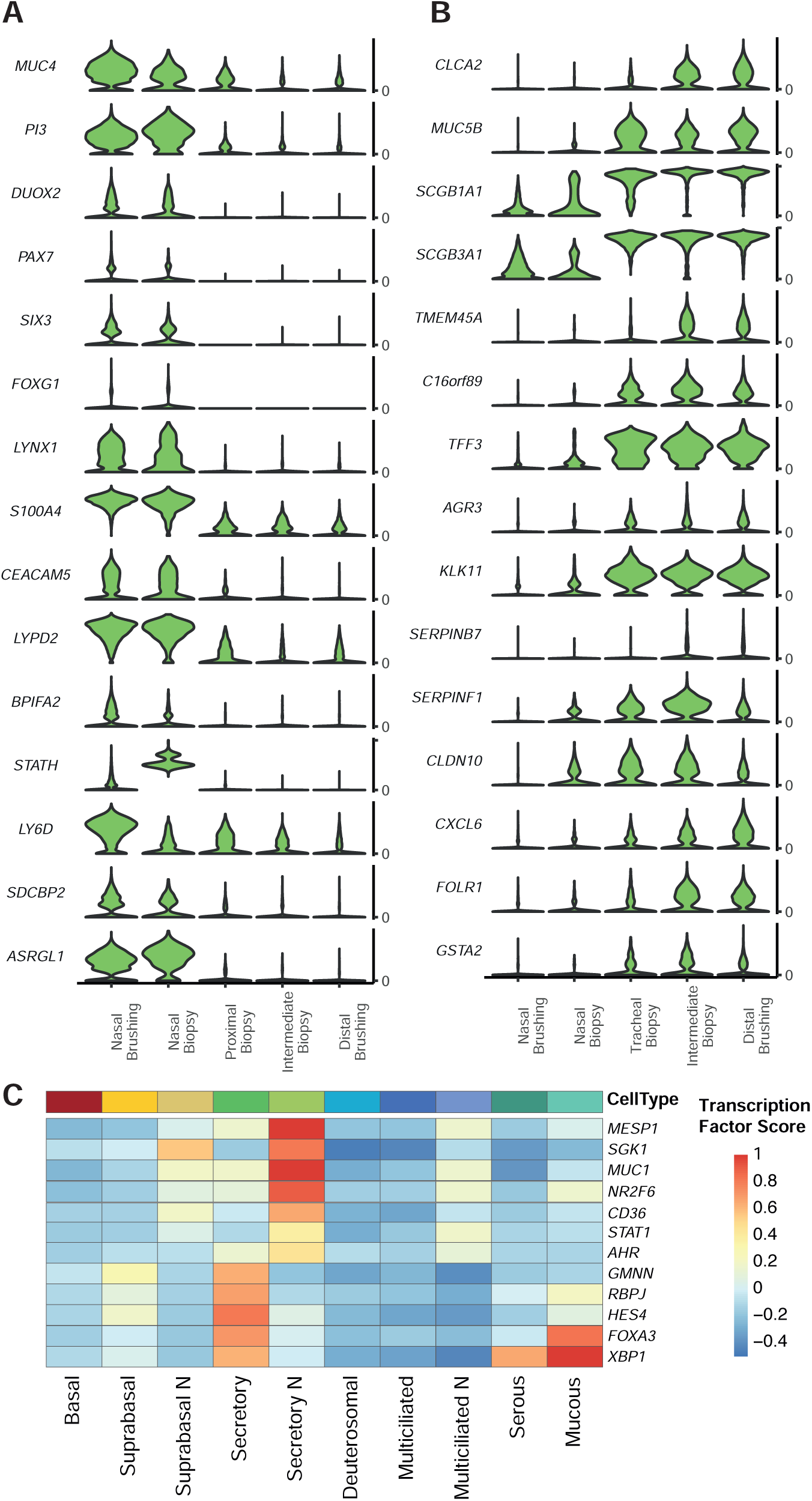
Nasal-Tracheobronchial specificities in gene expression in secretory cells. (A) Violin plot of up-regulated genes in nasal secretory cells (Secretory N). (B) Violin plot of up-regulated genes in tracheobronchial secretory cells. (C) Heatmap of cell type-specific regulatory unit activity score.

**Figure E10.**
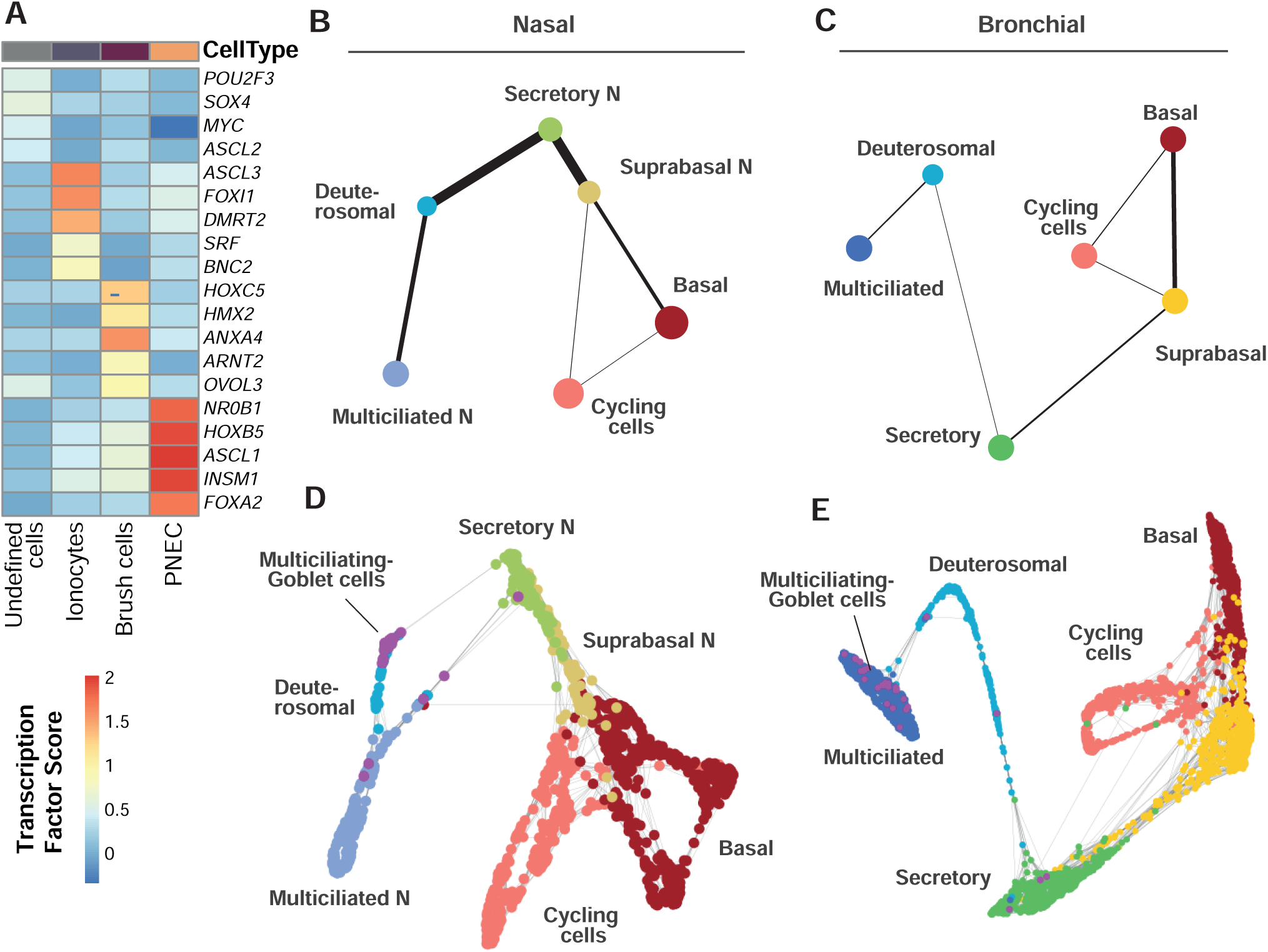
Rare cells detailed description. **(A)** Heatmap of specific regulatory unit activity score of rare cell type. (B-C) PAGA representations of airway epithelial cell lineages in nasal (B) and tracheobronchial epithelium (C). (D-E) Force atlas embedding of the inferred trajectory with superimposed mucous-multiciliated cells (purple) in nasal (D) and tracheobronchial epithelium (E).

**Figure E11.**
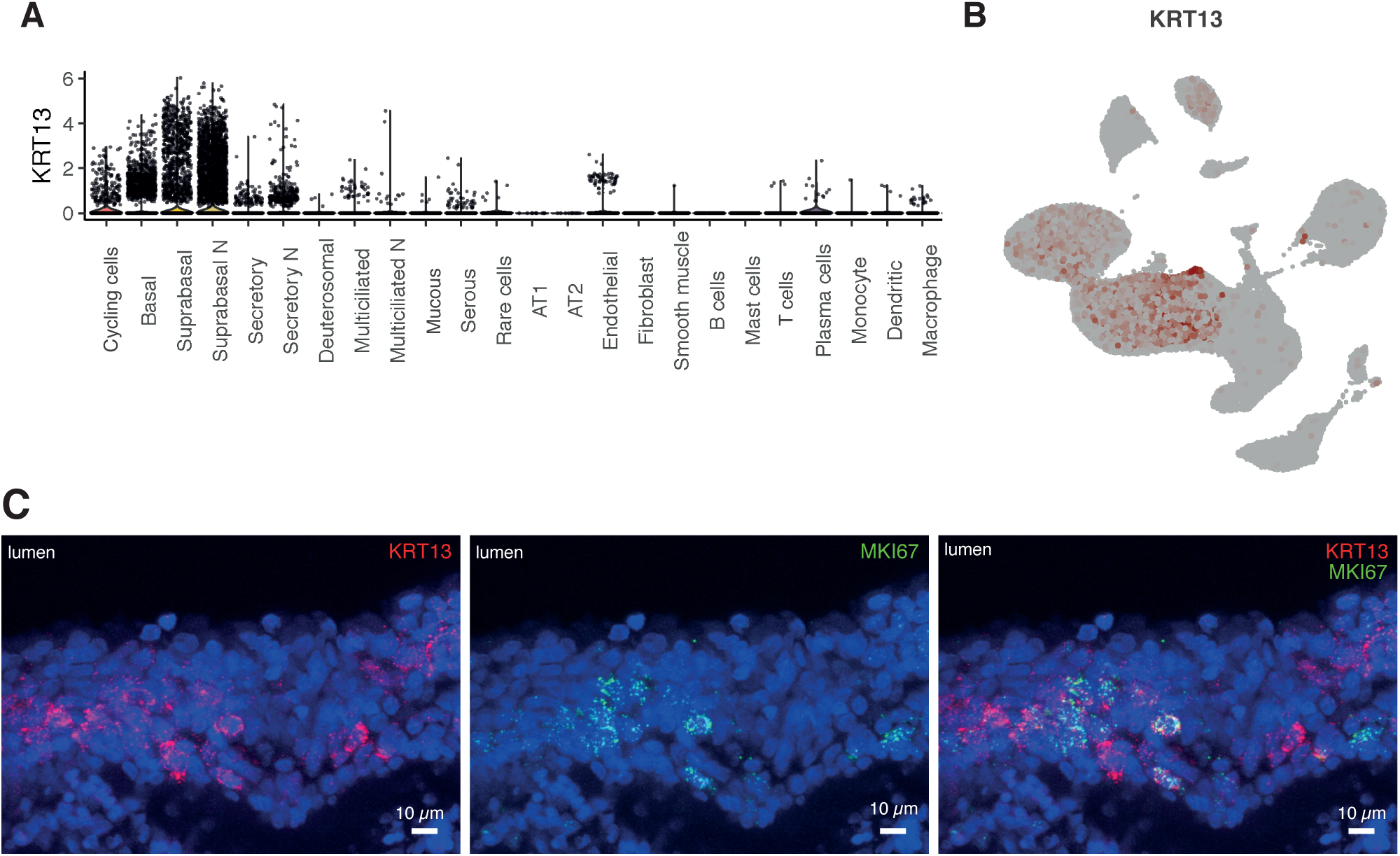
Identification and characterization of *KRT13*-positive cells. (A) Violin plot of the expression of *KRT13* by cell type (B) UMAP representation colored by the expression of *KRT13* and *KRT4* respectively. (C) RNAscope of KRT13 and MKI67 in nasal epithelial tissue.

